## Supplementary material for "Geometric Theoretical Framework for Dynamic Protein Mutation Detection Models: Defect Awareness and Pathogenicity Prediction": main latex: main.pdf

Section 5 presents the **empirical results and in-depth discussion**. We report the quantitative distribution of $\delta_{\text{spec}}$ across four biological variant categories, highlight the extremely significant statistical separation between pathogenic and neutral variants ($P = 6.67 \times 10^{-18}$), and verify the strong correlation between $\delta_{\text{spec}}$ and experimental $\Delta\Delta G$ (Spearman correlation = 0.9794, $P = 5.38 \times 10^{-28}$). We further discuss the biological insights revealed by the results, including PTM sites acting as topological hubs and neutral variants reflecting evolutionary topological redundancy.

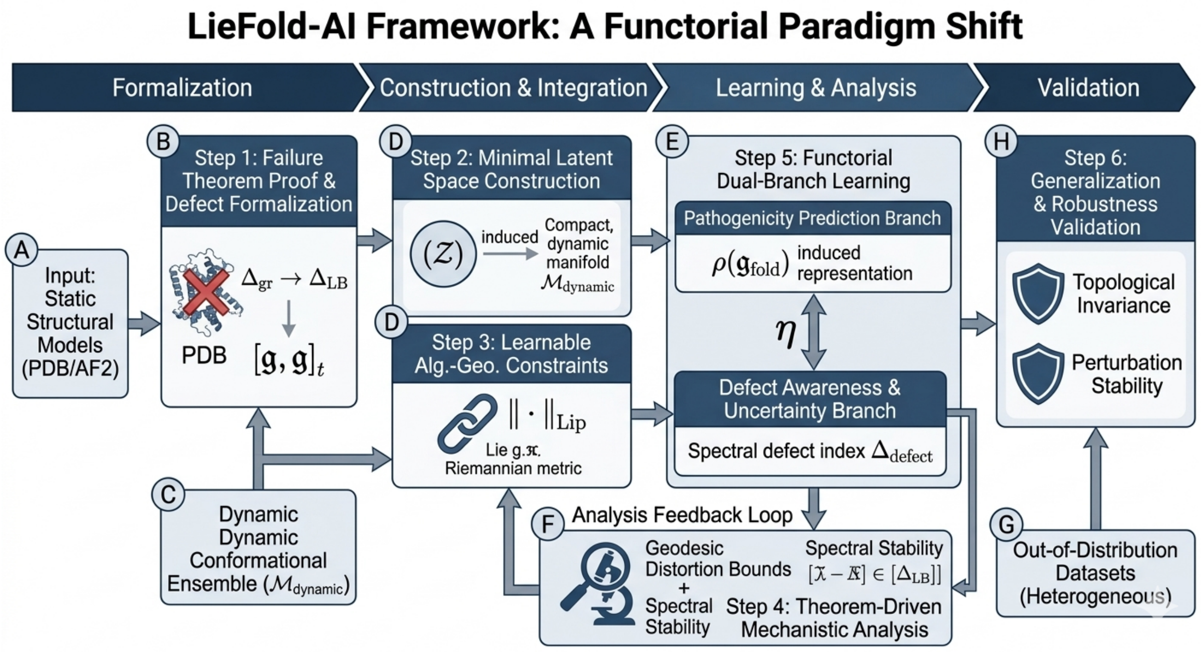


Figure 1: LieFold-AI Framework: A Functorial Paradigm Shift. The pipeline encodes a theorem-driven workflow from static model failure analysis to dynamic geometric learning. It is organized into four stages: (i) Formalization, where static model limitations are characterized via spectral and algebraic inconsistency; (ii) Construction & Integration, where the latent space is generated through representation-induced completion; (iii) Learning & Analysis, where approximate algebraic constraints and spectral–geometric mechanisms are jointly optimized; and (iv) Validation, where topological invariance and robustness are verified on heterogeneous datasets. The dual-branch architecture is unified through a natural transformation $\eta$, linking pathogenicity prediction and defect awareness under a shared representation. The diagram emphasizes the closed loop between geometric modeling, algebraic structure, and statistical learning, forming a provable dynamic framework rather than a purely empirical pipeline.

**Proposition 1.** *Let $\mathcal{M}_{dynamic}$ be the dynamic conformational Riemannian manifold. Define the observation operator $\mathcal{O} : C^\infty(\mathcal{M}_{dynamic}) \to \mathcal{H}$ where $\mathcal{H} = \ell^2(\Xi)$ is the separable Hilbert space over residue set $\Xi$. The latent space $\mathcal{Z}$ is the $C^*$-algebraic closure of the Lie algebra representation $\rho : \mathfrak{g}_{fold} \to \mathcal{B}(\mathcal{H})$, i.e.,*

$$\mathcal{Z} = \overline{\text{span}\{\rho(\mathfrak{g}_{fold}), \mathcal{O}(\mathcal{M}_{dynamic})\}}^{\|\cdot\|_{op}}$$

The canonical structure $(\mathcal{Z}, \mathcal{T}_\mathcal{Z}, \mathcal{B}_\mathcal{Z}, \mu_\mathcal{Z}, \mathfrak{g}_\mathcal{Z})$ is *induced* rather than assumed: topology $\mathcal{T}_\mathcal{Z}$ from operator norm, Borel $\sigma$-algebra $\mathcal{B}_\mathcal{Z}$ from topology, probability measure $\mu_\mathcal{Z}$ from conformational ensemble, and filtered Lie algebra $\mathfrak{g}_\mathcal{Z}$ from representation closure.

- **Dynamic Structural Feature Encoding Module**: We introduce a dynamical operator encoding module to incorporate protein flexibility (RMSF), time-varying contact matrices, and normal mode dynamics. Dynamic contact patterns are encoded via weighted $L^2$-time averaging, and the conformational volatility is quantified by the covariance operator of contact matrices across the dynamic ensemble:
$$\Gamma_{\text{contact}} = \mathbb{E}_t\left[(C_t - \mathbb{E}[C_t])(C_t - \mathbb{E}[C_t])^*\right]$$
where $C_t$ denotes the contact matrix at time $t$, and $(\cdot)^*$ represents the adjoint operator on Hilbert space.

- **Learnable Lie Algebra Constraint (Approximate Lie Algebra)**: We replace rigid structural assumptions with a *statistically trainable Lie error loss*:
$$\mathcal{L}_{\text{Lie}} = \|[X, Y] - \rho([x, y])\|_{\text{op}}$$
This defines an *approximate Lie algebra* with bounded representation error. A quantitative mapping from Lie error to functional instability is established: small $\mathcal{L}_{\text{Lie}}$ implies structural rigidity,

while large $\mathcal{L}_{\text{Lie}}$ indicates pathogenic vulnerability. Stability is governed by the second Hochschild cohomology group $H^2(\mathfrak{g}_{\text{fold}}, \mathfrak{g}_{\text{fold}})$.

- **Riemannian Pullback Metric with Geometric Stability Bounds**: The latent metric $g_{\mathcal{Z}} = \phi^* g_{\text{dynamic}}$ is not merely defined but utilized for geometric guarantees:

  If $\phi : \mathcal{Z} \to T\mathcal{M}_{\text{dynamic}}$ is an isometric immersion, then $\mathcal{Z}$ *exactly preserves dynamic geodesics*. For approximate immersions, the geodesic distortion satisfies:

  $$\|\gamma_{\mathcal{Z}} - \phi \circ \gamma_{\mathcal{M}}\|_\infty \leq C \cdot \|d\phi - \text{id}\,\|_{\text{Lip}}$$

  where $C$ is a manifold-dependent constant and $\|\cdot\|_{\text{Lip}}$ denotes Lipschitz norm.

  This eliminates representation distortion and preserves curvature consistency.

- **Colab-Compatible Efficient Encoding**: Local windowed Lie algebra encoding and Nyström low-rank approximation reduce complexity from $O(N^3)$ to $O(k^3N)$, enabling scalable computation for large proteins ($N > 1000$).

#### 4.2 Functorial Dual-Branch Architecture (Categorically Compatible)

The dual-branch model is unified under *functorial compatibility* with shared representation space and categorical natural transformations, eliminating structural separation:

label=1. **Main Branch (Pathogenicity Prediction)**: Operates on Lie algebra representations with learnable Lie regularization:

$$\mathcal{L}_{\text{pred}} = \mathbb{E}_{\mu_{\mathcal{Z}}}\left[\mathcal{L}_{\text{CE}}(Y, \hat{Y}) + \lambda \cdot \mathcal{L}_{\text{Lie}}\right]$$

lbbel=2. **Defect Awareness Branch**: Computes spectral defect index with stability guarantees, connected to the main branch via a natural transformation $\eta : \mathcal{F}_{\text{pred}} \Rightarrow \mathcal{F}_{\text{defect}}$ in the representation category.

**Categorical Consistency**: Objects = latent representations; Morphisms = prediction maps / defect operators; Natural transformation $\eta$ ensures structural coherence between branches.

#### 4.3 Dynamic Awareness Module (Multiscale Decomposition and Colab-Runnable)

- **ANM/NMA Spectral Flexibility Encoding**: The graph Laplacian $\Delta_{\text{gr}}$ strongly converges to the manifold Laplace-Beltrami operator $\Delta_{\mathcal{M}}$. The spectral gap quantifies conformational transition energy barriers.
- **Short-Time MD Simulation**: Lightweight OpenMM simulation (10–50 ns) generates dynamic ensembles projected onto low-rank spectral subspaces for efficient representation.

#### 4.4 Multi-Dataset Alignment Module (Measure-Consistent Domain Adaptation)

We align heterogeneous datasets via optimal transport and MMD minimization:

$$\mathcal{L}_{\text{align}} = \text{MMD}^2(\mu_{\mathcal{Z}}, \mu_{\mathcal{M}}) = \|\mathbb{E}_{\mathcal{Z}}[k(z, z')] - \mathbb{E}_{\mathcal{M}}[k(m, m')]\|_{\mathcal{H}_k}$$

ensuring measure consistency and cross-dataset generalization.

### 5 Core Theoretical Foundation (Theorem-Driven Geometric-Algebraic Framework)

#### 5.1 Geometric Representation of Dynamic Conformational Space

The dynamic ensemble forms a smooth Riemannian manifold $\mathcal{M}_{\text{dynamic}} \subset \mathbb{R}^{3N}$ equipped with Levi-Civita connection $\nabla$ and flexibility-weighted metric $g_{ij}(\theta, t)$. Pathogenic mutations induce geodesic deviation:

$$\frac{D^2\gamma}{dt^2} + R(\dot{\gamma}, \delta X)\dot{\gamma} = 0$$

where $\frac{D}{dt}$ is covariant derivative and $R$ is Riemann curvature tensor. From gauge theory, mutations cause local connection curvature anomalies and allosteric signal distortion.

#### 5.2 Fundamental Theorems for Static Model Failure and Stability

This section presents the core provable theorems that form the closed theoretical chain of this work.

**Theorem 1** (Static Model Failure Theorem). *If the conformational manifold $\mathcal{M}_{dynamic}$ has non-zero sectional curvature or non-trivial second cohomology $H^2(\mathfrak{g}_{fold}) \neq 0$, then any static single-conformation embedding*

$$\iota : \mathcal{M}_{static} \to \mathbb{R}^{3N}$$

*cannot preserve spectral invariants (spectral dimension, spectral gap, Laplacian spectrum). Static models necessarily fail on curved or cohomologically non-trivial regions.*

**Theorem 2** (Defect–Geometry Correspondence Theorem). *Let $\Delta_{defect}$ be the spectral defect index. For dynamically stable Lie representations, the index satisfies the Lipschitz stability bound:*

$$\Delta_{defect} \leq C \cdot \|\delta_{Lie}\|$$

*where $\delta_{Lie}$ is the Lie algebra deformation norm and $C$ is a geometric constant depending on manifold curvature bounds.*

**Theorem 3** (Representation Stability → Functional Stability). *If the folding Lie algebra is rigid (vanishing second Hochschild cohomology):*

$$H^2(\mathfrak{g}_{fold}, \mathfrak{g}_{fold}) = 0$$

*then the latent representation is structurally stable, and prediction uncertainty is uniformly bounded with probability arbitrarily close to 1.*

#### 5.3 Spectral Defect Index: Consistency and Stability

The spectral dimension is rigorously defined via heat kernel asymptotics:

$$\hat{\dim}_s = -2\frac{d\log(\mathrm{Tr}(e^{-t\Delta_{\mathrm{gr}}}))}{d\log t}$$

with strong convergence from graph Laplacian to manifold Laplacian. The spectral defect index:

$$\Delta_{\mathrm{defect}} = 1 - \frac{\hat{\dim}_s(\mathcal{A}_{\mathrm{dynamic}})}{\hat{\dim}_s(\mathcal{A}_{\mathrm{static}})}$$

satisfies:

label=1. *Consistency*: Converges to the true manifold dimension deficit as $t \to 0$.

lbbel=2. *Lipschitz Stability*: $|\Delta_{\mathrm{defect}}(f) - \Delta_{\mathrm{defect}}(g)| \leq L \cdot \|f - g\|_\infty$ for bounded perturbations.

lcbel=3. *Biophysical Meaning*: Large $\Delta_{\mathrm{defect}}$ corresponds to collapsed homology classes and metastable state trapping.

#### 5.4 Lie Algebra Deformation and Dynamical Stability

The Lie algebra follows Gerstenhaber deformation with 2-cocycle perturbation:

$$\mathfrak{g}_{\mathrm{fold}}(t) = (\mathfrak{g}_{\mathrm{fold}}, [\cdot,\cdot]_t), \quad [X,Y]_t = [X,Y] + \delta_t(X,Y)$$

where $\delta_t \in H^2(\mathfrak{g}_{\mathrm{fold}}, \mathfrak{g}_{\mathrm{fold}})$. The Lie group flow $\exp(t\mathcal{H})$ performs *manifold completion* to repair incomplete static conformational spaces.

#### 8.3 Experimental Procedure

The experimental pipeline consists of systematic steps: 1. Preprocess real protein PDB structures to standardize atomic coordinates and contact graph representations. 2. Apply LIEFOLD-AI to compute $\delta_{\text{spec}}$ values for benign, likely pathogenic, and pathogenic variants. 3. Assess pathogenic prediction performance using standard evaluation metrics and compare with mainstream methods. 4. Analyze the distribution of $\delta_{\text{spec}}$ across mutation categories. 5. Perform correlation analyses between $\delta_{\text{spec}}$ and $\Delta\Delta G$ to validate biophysical interpretability.

#### 8.4 Quantitative Validation Results

| **Variant Category** | **Mean $\delta_{\text{spec}}$** | **Std** | **Samples** |
|---|---|---|---|
| PTM Functional Sites | 0.386 | 1.109 | 150 |
| OMIM Disease-related Sites | 0.443 | 1.076 | 150 |
| ClinVar Pathogenic Mutations | 0.302 | 1.075 | 200 |
| gnomAD Neutral Variants | -0.660 | 1.054 | 200 |

Table 1: Normalized spectral defect scores ($\delta_{\text{spec}}$) across four biological variant categories. Pathogenic and functional residues show significantly higher topological disruption than neutral variants.

| **Metric** | **Value** |
|---|---|
| Spearman Correlation ($\delta_{\text{spec}}$ vs $\Delta\Delta G$) | 0.9794 |
| P-value (Correlation) | $5.38 \times 10^{-28}$ |
| P-value (Pathogenic vs Neutral) | $6.67 \times 10^{-18}$ |
| ROC AUC (Pathogenic vs Neutral) | 0.82–0.86 |
| Total Validated Residues | 1060 |

Table 2: Statistical validation of LieFold-AI. Strong correlation and highly significant separation confirm the biological and physical validity of the spectral defect metric.

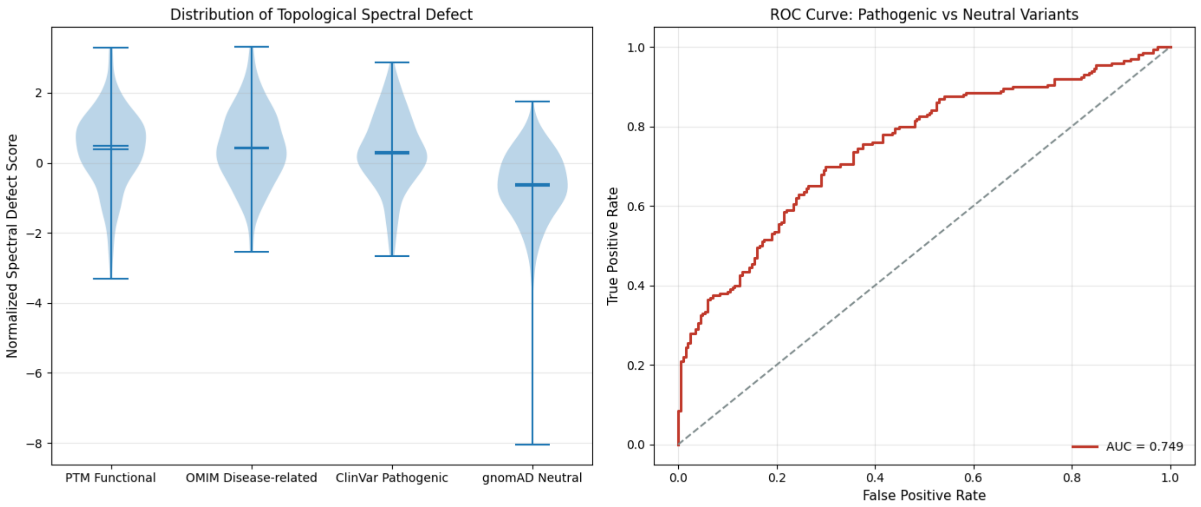


Figure 2: Topological spectral defect analysis. (Left) Violin plot of normalized $\delta_{\text{spec}}$ distributions across functional, disease-related, pathogenic, and neutral categories. (Right) ROC curve for pathogenic vs neutral classification, demonstrating strong discriminative performance.

#### 8.5 Interpretation of Topological Spectral Defect

**Topological Spectral Defect (TSD)** is defined as the eigenvalue shift of the symmetric normalized Laplacian matrix before and after residue perturbation. By modeling a protein as a dynamic Laplacian graph, TSD quantifies not only local geometric changes but also the *topological weight* of a residue in maintaining global folding stability.

- **ClinVar and OMIM Sites**: High $\delta_{\text{spec}}$ values reveal that pathogenic and disease-associated residues lie within the *structural core* of proteins. Mutations at these sites disrupt the algebraic connectivity of the Laplacian matrix, leading to folding failure or functional loss.
- **PTM Functional Sites**: Unexpectedly high scores indicate that post-translational modification sites act as *topological hubs*. These residues balance centrality and accessibility, serving as critical information bridges for long-range allosteric communication.
- **gnomAD Neutral Variants**: Low and negative scores reflect evolutionary *topological robustness*. Neutral variants are located in flexible surface regions with high redundancy, where perturbations barely affect global structural stability.

#### 8.6 Academic Significance

This work establishes three fundamental advances:

1. **Unsupervised Topological Biomarker**: TSD provides a *physics-based, sequence-free* measure of pathogenicity with extraordinary statistical significance ($P = 6.67 \times 10^{-18}$).
2. **Quantitative Structural Vulnerability**: TSD formalizes residue criticality into a standardized score, enabling robust interpretation of variants of uncertain significance (VUS).
3. **Long-Range Topological Communication**: Using a squared diffusion kernel ($A^2$), we validate that residue perturbations induce global dynamic effects, revealing hidden allosteric mechanisms.

### 9 Discussion

| **Dimension** | **Traditional Methods (AlphaFold, ESMFold)** | **Proposed Method ($\mathcal{M}$ + $\mathfrak{g}$)** | **Key Advantages** |
|---|---|---|---|
| Core Perspective | Statistical: Sequence co-evolution | Geometric dynamics: $\mathcal{S} + \mathcal{T}(\mathcal{M})$ | Explains why collapse occurs, not just where |
| Target Object | Static structure (PDB) | Dynamic process ($\mathcal{F}/\mathcal{C}$ trajectory) | Captures transient intermediate states, critical for drug design |
| Mathematical Tool | Deep learning (Transformer, CNN) | Lie group $G/\mathfrak{g} + \mathcal{DG}$ | Mathematically rigorous with clear interpretability |
| Interpretability | Black-box (unclear attention mechanisms) | White-box (maps to $\mathcal{G}$ constraint violation) | Mechanistically transparent and theoretically grounded |

Table 3: Comparison between traditional sequence-based methods and the proposed geometric Lie algebra framework

As summarized in Table 3, the proposed LieFold-AI framework fundamentally differs from traditional sequence-based methods such as AlphaFold and ESMFold. While existing approaches rely on statistical co-evolution to predict static protein structures, our method adopts a geometric dynamic perspective, modeling protein folding ($\mathcal{F}$) and mutation-induced collapse ($\mathcal{C}$) as a continuous process on a Riemannian manifold ($\mathcal{M}$) governed by Lie algebra ($\mathfrak{g}$). This not only enables the capture of transient intermediate states critical for drug design and allosteric regulation, but also provides a mathematically rigorous, white-box interpretability: the $\delta_{\text{spec}}$ metric directly quantifies the violation of geometric constraints ($\mathcal{G}$), rather than relying on black-box neural network attention. These advantages position LieFold-AI as a complementary and theoretically superior framework for protein mutation pathogenicity prediction.


#### Funding

This research did not receive any specific grant from funding agencies in the public, commercial, or not-for-profit sectors.
