## Supplementary material for "Geometric Theoretical Framework for Dynamic Protein Mutation Detection Models: Defect Awareness and Pathogenicity Prediction": main latex: main.tex

\documentclass{article}

% 页面布局
\usepackage[a4paper, margin=1in]{geometry}

% 数学环境
\usepackage{amsmath, amssymb, amsthm}

% 表格
\usepackage{tabularx}

% 列表（兼容 enumerate）
\usepackage{enumerate}

% 图片
\usepackage{graphicx}

% 超链接（建议最后加载）
\usepackage{hyperref}

% =========================
% 定理环境
% =========================
\theoremstyle{plain}
\newtheorem{theorem}{Theorem}
\newtheorem{lemma}{Lemma}
\newtheorem{proposition}{Proposition}
\newtheorem{corollary}{Corollary}[proposition]

\theoremstyle{definition}
\newtheorem{definition}{Definition}


% =========================
% 数学算子（推荐用 \DeclareMathOperator 更规范）
% =========================
\DeclareMathOperator{\Tr}{Tr}
\DeclareMathOperator{\diag}{diag}
\DeclareMathOperator{\Var}{Var}
\DeclareMathOperator{\ad}{ad}
\DeclareMathOperator{\diam}{diam}
% 标题
\title{Geometric Theoretical Framework for Dynamic Protein Mutation Detection Models: Defect Awareness and Pathogenicity Prediction}


\author{
	Haijian Shao\\
	School of Computer,\\
	Jiangsu University of Science and Technology, Zhenjiang 212003, China\\
	\small 
%	\and
%	Daze Lu, Yingtao Jiang\\
%	Department of Electrical and Computer Engineering,\\
%	University of Nevada, Las Vegas, USA
}
\date{}

%% 单栏全局设置
%\onecolumn
\begin{document}
	\maketitle
	
\begin{abstract}
	Traditional protein mutation detection and pathogenicity prediction pipelines rely on static single-conformation structural modeling, inherently ignoring conformational flexibility, dynamic ensemble evolution, and the underlying manifold geometry of protein dynamics. This induces systematic detection failures in flexible regions, allosteric sites, and metastable functional domains, yet lacks a rigorous mathematical characterization of such failure mechanisms. In this work, we establish a theorem-driven geometric-algebraic framework for dynamic protein mutation modeling. Starting from a dynamic conformational Riemannian manifold, we construct the latent representation space via representation-induced completion of operator-valued observations, rather than pre-assumed embedding structures. Within this setting, algebraic constraints are not imposed axiomatically but relaxed into learnable approximate Lie algebra regularization, enabling statistical verification of structural consistency.
	By integrating Levi-Civita connection, geodesic deviation, and heat kernel asymptotics, we introduce a Lipschitz-stable topological spectral defect (TSD, $\delta_{\text{spec}}$) index that quantifies the intrinsic inconsistency between static representations and dynamic geometric invariants, linking it to curvature-induced instability and Lie algebra deformation. Under a functorial compatibility principle, we design a dual-branch architecture for pathogenicity prediction and defect awareness, realized via local Lie algebra encoding and low-rank spectral approximation.
	On multi-source datasets (108 curated PDB structures, 1060 validated residues from ClinVar, DMS, MaveDB, and gnomAD), we establish three fundamental theorems and validate key findings: TSD effectively distinguishes pathogenic/functional variants (PTM: $\mu=0.386$, OMIM: $\mu=0.443$, ClinVar: $\mu=0.302$) from neutral ones (gnomAD: $\mu=-0.660$) with high significance ($P=6.67 \times 10^{-18}$) and strong classification performance (AUC=0.82–0.86), while correlating strongly with protein stability ($\Delta\Delta G$, Spearman=0.9794, $P=5.38 \times 10^{-28}$). TSD further reveals PTM sites as topological hubs and neutral variants as evolutionary topological redundancy, enabling a paradigm shift from sequence alignment to geometric dynamics and providing a physics-based biomarker for variants of uncertain significance (VUS). These results upgrade protein mutation modeling from empirical static prediction to provable dynamic mechanism analysis. The source code of this work is publicly available at \url{https://github.com/Harmenlv/LieFold-AI/tree/main}.
\end{abstract}

\section{Introduction}

Understanding the functional and pathogenic impact of single amino acid variants (SAVs) is essential for clinical variant interpretation, precision medicine, and drug discovery. 
Amino acid substitutions can disrupt protein folding, stability, interaction, and allosteric communication, leading to various genetic diseases and complex disorders. 
Publicly available databases such as ClinVar, OMIM, and gnomAD have accumulated large-scale curated variant annotations, yet the biophysical and structural mechanisms underlying pathogenicity remain incompletely understood. 
Many variants of uncertain significance (VUS) lack reliable interpretability due to the absence of physically grounded predictive models, creating a critical gap between genomic data and clinical decision-making.

Traditional computational methods for variant effect prediction rely heavily on sequence homology, evolutionary conservation, and statistical feature engineering, including widely used tools such as SIFT, PolyPhen-2, and CADD. 
Although these methods achieve decent performance in many scenarios, they lack explicit structural and physical interpretability. 
With the rapid development of deep learning, advanced models such as AlphaMissense and structure-based predictors have improved prediction accuracy by integrating protein language models and static structural information~\cite{jumper2021highly}. 
Geometric deep learning has further enabled powerful representations for structure-based drug design and protein analysis~\cite{isert2023structure}. 
However, most of these models remain black-box systems, whose decisions cannot be easily traced to biophysical principles. 
Recent self-supervised geometric learning methods have attempted to improve interpretability for mutation-induced stability change prediction~\cite{li2024prediction}, and state-of-the-art pathogenicity predictors such as Rhapsody-2 provide mechanistic evaluation of missense variants~\cite{banerjee2025accurate}. 
Despite these advances, nearly all mainstream methods treat proteins as static or quasi-static structures, ignoring conformational flexibility, dynamic ensemble behavior, and the global topological organization of protein folds. 
As a result, such methods fail to capture long-range structural effects, allosteric perturbations, and metastable state transitions, which are central to mutation-induced functional collapse.

Geometric and topological approaches have emerged as promising tools to model protein structure and dynamics from first principles. 
Riemannian geometry has been successfully applied to protein structure determination~\cite{wei2020protein} and the efficient analysis of protein dynamics data~\cite{diepeveen2024riemannian}. 
Classical and modern geometric analyses have revealed the fundamental role of topological organization in protein folding~\cite{simmons2008protein, begun2025local}. 
In particular, local topological transitions and perestroikas directly govern folding dynamics and structural robustness~\cite{begun2025local}, while hydrophobic collapse represents a key biophysical signature of structural integrity~\cite{walther2007signatures}. 
The mathematical foundation of spectral geometry~\cite{atiyah1975spectral} provides a rigorous basis for quantifying geometric invariance and perturbation effects, which has not yet been fully exploited for variant pathogenicity assessment.

To address these limitations, this study introduces a unified geometric theoretical framework for dynamic protein mutation detection and pathogenicity prediction, centered on the notion of \textit{defect awareness} and topological spectral analysis. 
In contrast to conventional methods that view proteins as static sequences or fixed conformations, we model a protein as a dynamic Riemannian manifold and a continuous Laplacian graph, where mutation-induced damage is quantified by a theoretically grounded measure called \textit{Topological Spectral Defect} (TSD, $\delta_{\text{spec}}$). 
This framework is rooted in Lie algebra, graph dynamics, and differential geometry, enabling a white-box characterization of how local perturbations propagate globally and disrupt structural stability. 
The proposed framework shifts the paradigm from empirical sequence-based prediction to principle-driven geometric analysis, where pathogenicity is directly explained by topological constraint violation rather than statistical correlation.

The contributions of this work are threefold. 
First, we establish a formal geometric foundation for protein variant analysis by constructing a dynamic conformational manifold with learnable Lie algebra regularization, enabling Lipschitz-stable quantification of structural deformation. 
Second, we propose the Topological Spectral Defect (TSD) index, enhanced by a long-range $A^2$ diffusion kernel and local Z-score normalization, which reliably distinguishes pathogenic, disease-related, functional, and neutral variants in an unsupervised manner. 
Third, we validate the framework extensively on real-world datasets, including 108 high-quality PDB structures, ClinVar pathogenic mutations, gnomAD neutral variants, OMIM disease-associated sites, and PTM functional sites, achieving highly significant separation ($P = 6.67 \times 10^{-18}$) and strong classification performance (AUC = 0.82–0.86). 
The index also exhibits an extremely strong correlation with experimental protein stability ($\Delta\Delta G$), confirming its biophysical validity.

\textbf{Paper organization:}
To present the research in a logically coherent and systematic manner, the structure of this paper is arranged as follows, following the progression from theoretical construction to experimental validation and practical implications. 

First, Section 2 establishes the \textbf{Geometric Theoretical Framework} for dynamic protein mutation modeling. We formally define the protein as a dynamic Laplacian graph on a Riemannian manifold, derive the mathematical foundation of the Topological Spectral Defect (TSD, $\delta_{\text{spec}}$) index, and elaborate on the connection between TSD, Lie algebra deformation, and curvature-induced structural instability—laying the theoretical basis for defect awareness.

Building upon this theoretical foundation, Section 3 details the \textbf{LieFold-AI methodology and implementation}. We describe the complete pipeline, including the integration of the $A^2$ long-range diffusion kernel for capturing cross-domain topological communication, the symmetric Laplacian decomposition for stable spectral computation, and the local background Z-score normalization strategy to eliminate protein size bias and ensure generalization across diverse protein folds. Notably, this pipeline adopts an unsupervised learning paradigm, avoiding reliance on large-scale labeled mutation data.

Section 4 outlines the \textbf{experimental setup and dataset details} to validate the proposed framework. We introduce the large-scale curated datasets used in this study, including 108 high-quality real PDB structures, ClinVar pathogenic mutations, gnomAD neutral variants, OMIM disease-associated sites, PTM functional sites, and Deep Mutational Scanning (DMS) $\Delta\Delta G$ data—ensuring rigorous and comprehensive validation of both predictive performance and biophysical relevance.

Section 5 presents the \textbf{empirical results and in-depth discussion}. We report the quantitative distribution of $\delta_{\text{spec}}$ across four biological variant categories, highlight the extremely significant statistical separation between pathogenic and neutral variants ($P = 6.67 \times 10^{-18}$), and verify the strong correlation between $\delta_{\text{spec}}$ and experimental $\Delta\Delta G$ (Spearman correlation = 0.9794, $P = 5.38 \times 10^{-28}$). We further discuss the biological insights revealed by the results, including PTM sites acting as topological hubs and neutral variants reflecting evolutionary topological redundancy.

Finally, Section 6 concludes the paper by summarizing the core contributions of the LieFold-AI framework to unsupervised mutation pathogenicity prediction and geometric protein modeling. We also outline future research directions, including the integration of structural dynamics into clinical variant interpretation (e.g., for variants of uncertain significance, VUS) and the extension of the framework to allosteric mutation analysis. The source code of this work is publicly available at \url{https://github.com/Harmenlv/LieFold-AI/tree/main}.

	
\section{Research Positioning and Core Logic}
\subsection{Review of Previous Work}
Classical studies are confined to the paradigm of static protein folding geometry, taking single rigid conformations (native PDB structures or AlphaFold-predicted structural models) as the sole carrier of structural information. The research adopts standard benchmark datasets including ClinVar, DMS, 102 high-resolution native PDB structures, and 5,000 homology-modeled simulated mutation structures. The core objective is restricted to the static supervised prediction of amino acid mutational effects and pathogenicity classification, lacking rigorous mathematical characterization of model detection defects, and completely ignoring the dynamic conformational ensemble and intrinsic flexibility of proteins, resulting in the absence of dynamic mechanism analysis.

\subsection{Positioning and Core Innovation of the New Work}
This work abandons the repetitive construction of traditional static prediction pipelines, and centers on the systematic detection defects inherent to amino acid mutation sites under static geometric modeling, realizing a fundamental paradigm shift from static single-conformation analysis to dynamic conformational ensemble learning. The research content has no conceptual overlap with existing work, and the theoretical depth and methodological rigor are comprehensively upgraded. Meanwhile, we expand multi-source high-throughput heterogeneous datasets, and all experimental protocols are reproducible on Colab, which greatly strengthens the theoretical persuasiveness and empirical reliability of the paper.

\textbf{Core Innovation (One-Sentence Summary)}:
Aiming at the systematic detection defects of static protein geometric models on ClinVar-annotated pathogenic sites, we construct a unified functorial geometric learning pipeline rooted in functional analysis and Lie theory, rigorously characterize failure modes via geometric invariance over large-scale multi-source datasets, incorporate dynamic conformational features encoded by Lie algebra representations and Riemannian geometry, and build an enhanced dual-branch model integrating pathogenicity prediction and intrinsic defect awareness, ultimately realizing the paradigm shift from static structural prediction to dynamic conformational mechanism analysis.

\section{Problem Statement and Research Methodology}
\subsection{Core Problem: Intrinsic Limitations of Static Models}
Traditional static structural models are restricted to the analysis of isolated protein conformations, which fundamentally ignores the non-rigid flexibility, continuous conformational fluctuations, and dynamic contact evolution of proteins in physiological environments. This inherent limitation induces systematic detection failures on specific amino acid sites (e.g., intrinsically flexible regions, time-varying dynamic contact interfaces, allosteric functional sites). Such models cannot capture the mutation-induced conformational dynamics encoded in the infinite-dimensional conformational space, nor can they reveal the underlying biophysical pathogenic mechanisms; more critically, they lack the mathematical self-awareness of their own prediction defects due to the incompleteness of static geometric encoding and broken manifold completeness.

\subsection{Complete Research Pipeline}
To clearly illustrate the practical implementation and theoretical structure of the LieFold-AI framework, we organize the workflow into a vector-style pipeline diagram (Fig.~\ref{fig:liefold-ai-framework}) together with six sequential and logically coupled steps. The diagram is not merely illustrative but encodes the theorem-driven construction of the framework, explicitly linking mathematical formalization, representation construction, and learning mechanisms into a unified pipeline. It follows a left-to-right flow and is divided into four stages: \textit{Formalization}, \textit{Construction \& Integration}, \textit{Learning \& Analysis}, and \textit{Validation}. Each stage corresponds to a necessary component in establishing a provable dynamic modeling framework for protein mutation detection.

\begin{enumerate}[label=\arabic*.]
	\item \textbf{Mathematical Problem Formalization}: We first establish a \emph{static model failure theorem} by showing that single-conformation representations cannot preserve dynamic spectral invariants. This is rigorously characterized via graph Laplacian convergence, heat kernel asymptotics, and Lie algebra deformation theory, providing a mathematical explanation for systematic prediction failures in flexible and allosteric regions.
	
	\item \textbf{Minimal Latent Space Construction}: Instead of predefining a full latent structure, we construct the latent space as a \textit{representation-induced closure} of operator-valued observations over the dynamic conformational manifold. This ensures that the latent space is generated from underlying geometric dynamics rather than imposed a priori, eliminating redundant structural assumptions.
	
	\item \textbf{Learnable Algebraic-Geometric Constraints}: Classical algebraic constraints are relaxed into trainable regularization terms with Lipschitz stability bounds. In particular, approximate Lie algebra consistency is enforced through operator-level penalties, making abstract structural constraints statistically verifiable during training.
	
	\item \textbf{Theorem-Driven Mechanistic Analysis}: Based on the constructed latent space, we derive spectral–geometric correspondence results, including defect stability bounds, geodesic distortion control, and representation stability. This step establishes a direct link between mutation-induced geometric variation and model prediction behavior.
	
	\item \textbf{Functorial Dual-Branch Learning}: We design a dual-branch architecture for joint pathogenicity prediction and defect awareness. The two branches are unified via a natural transformation $\eta$, ensuring consistent representation sharing under a functorial framework while preserving task-specific outputs.
	
	\item \textbf{Generalization and Robustness Validation}: We validate the framework on heterogeneous datasets (e.g., ClinVar, DMS, MaveDB), verifying topological invariance and perturbation stability under distribution shifts. This confirms that the proposed framework generalizes beyond training distributions and remains consistent with underlying geometric structures.
\end{enumerate}

\begin{figure*}[t]
	\centering
	\includegraphics[width=\textwidth]{pipeline.png}
	\caption{
		LieFold-AI Framework: A Functorial Paradigm Shift. 
		The pipeline encodes a theorem-driven workflow from static model failure analysis to dynamic geometric learning. 
		It is organized into four stages: (i) Formalization, where static model limitations are characterized via spectral and algebraic inconsistency; 
		(ii) Construction \& Integration, where the latent space is generated through representation-induced completion; 
		(iii) Learning \& Analysis, where approximate algebraic constraints and spectral–geometric mechanisms are jointly optimized; 
		and (iv) Validation, where topological invariance and robustness are verified on heterogeneous datasets. 
		The dual-branch architecture is unified through a natural transformation $\eta$, linking pathogenicity prediction and defect awareness under a shared representation. 
		The diagram emphasizes the closed loop between geometric modeling, algebraic structure, and statistical learning, forming a provable dynamic framework rather than a purely empirical pipeline.
	}
	\label{fig:liefold-ai-framework}
\end{figure*}

The pipeline reflects the central contribution of this work: transforming protein mutation modeling from an empirical static prediction paradigm into a theorem-driven dynamic geometric framework. By grounding representation learning in manifold geometry, Lie algebra structure, and spectral invariants, the proposed approach achieves a principled integration of mathematical rigor and practical applicability, providing a new foundation for interpretable and robust pathogenicity analysis.

\section{Model Improvement (Reconstruction and Upgrade Based on Original Framework)}
\subsection{Minimal Latent Space Construction (Representation-Induced Completion)}
We discard the predefined full-structure latent space paradigm and adopt a \textbf{minimal generative construction} following the principle: \textit{Latent space = representation-induced completion}. The construction follows a strict causal chain:

\begin{proposition}
	Let $\mathcal{M}_{\text{dynamic}}$ be the dynamic conformational Riemannian manifold. Define the observation operator $\mathcal{O}: C^\infty(\mathcal{M}_{\text{dynamic}}) \to \mathcal{H}$ where $\mathcal{H} = \ell^2(\Xi)$ is the separable Hilbert space over residue set $\Xi$. The latent space $\mathcal{Z}$ is the \textit{$C^*$-algebraic closure} of the Lie algebra representation $\rho: \mathfrak{g}_{\text{fold}} \to \mathcal{B}(\mathcal{H})$, i.e.,
	$$
	\mathcal{Z} = \overline{\operatorname{span}\left\{ \rho(\mathfrak{g}_{\text{fold}}), \mathcal{O}(\mathcal{M}_{\text{dynamic}}) \right\}}^{\|\cdot\|_{\text{op}}}
	$$
\end{proposition}

The canonical structure $(\mathcal{Z}, \mathcal{T}_\mathcal{Z}, \mathcal{B}_\mathcal{Z}, \mu_\mathcal{Z}, \mathfrak{g}_\mathcal{Z})$ is \textit{induced} rather than assumed: topology $\mathcal{T}_\mathcal{Z}$ from operator norm, Borel $\sigma$-algebra $\mathcal{B}_\mathcal{Z}$ from topology, probability measure $\mu_\mathcal{Z}$ from conformational ensemble, and filtered Lie algebra $\mathfrak{g}_\mathcal{Z}$ from representation closure.

\begin{itemize}
	\item \textbf{Dynamic Structural Feature Encoding Module}: We introduce a dynamical operator encoding module to incorporate protein flexibility (RMSF), time-varying contact matrices, and normal mode dynamics. Dynamic contact patterns are encoded via weighted $L^2$-time averaging, and the conformational volatility is quantified by the covariance operator of contact matrices across the dynamic ensemble:
	$$
	\Gamma_{\text{contact}} = \mathbb{E}_t\left[ (C_t - \mathbb{E}[C_t])(C_t - \mathbb{E}[C_t])^* \right]
	$$
	where $C_t$ denotes the contact matrix at time $t$, and $(\cdot)^*$ represents the adjoint operator on Hilbert space.
	
	\item \textbf{Learnable Lie Algebra Constraint (Approximate Lie Algebra)}: We replace rigid structural assumptions with a \textit{statistically trainable Lie error loss}:
	$$
	\mathcal{L}_{\text{Lie}} = \left\| [X,Y] - \rho([x,y]) \right\|_{\text{op}}
	$$
	This defines an \textit{approximate Lie algebra} with bounded representation error. A quantitative mapping from Lie error to functional instability is established: small $\mathcal{L}_{\text{Lie}}$ implies structural rigidity, while large $\mathcal{L}_{\text{Lie}}$ indicates pathogenic vulnerability. Stability is governed by the second Hochschild cohomology group $H^2(\mathfrak{g}_{\text{fold}}, \mathfrak{g}_{\text{fold}})$.
	
	\item \textbf{Riemannian Pullback Metric with Geometric Stability Bounds}: The latent metric $g_\mathcal{Z} = \phi^* g_{\text{dynamic}}$ is not merely defined but utilized for geometric guarantees:

		If $\phi: \mathcal{Z} \to T\mathcal{M}_{\text{dynamic}}$ is an isometric immersion, then $\mathcal{Z}$ \textit{exactly preserves dynamic geodesics}. For approximate immersions, the geodesic distortion satisfies:
		$$
		\|\gamma_\mathcal{Z} - \phi \circ \gamma_{\mathcal{M}}\|_\infty \leq C \cdot \|d\phi - \operatorname{id}\|_{\text{Lip}}
		$$
		where $C$ is a manifold-dependent constant and $\|\cdot\|_{\text{Lip}}$ denotes Lipschitz norm.


	This eliminates representation distortion and preserves curvature consistency.
	
	\item \textbf{Colab-Compatible Efficient Encoding}: Local windowed Lie algebra encoding and Nyström low-rank approximation reduce complexity from $O(N^3)$ to $O(k^3 N)$, enabling scalable computation for large proteins ($N>1000$).
\end{itemize}

\subsection{Functorial Dual-Branch Architecture (Categorically Compatible)}
The dual-branch model is unified under \textit{functorial compatibility} with shared representation space and categorical natural transformations, eliminating structural separation:

\begin{enumerate}[label=\arabic*.]
	\item \textbf{Main Branch (Pathogenicity Prediction)}: Operates on Lie algebra representations with learnable Lie regularization:
	$$
	\mathcal{L}_{\text{pred}} = \mathbb{E}_{\mu_\mathcal{Z}}\left[ \mathcal{L}_{\text{CE}}(Y, \hat{Y}) + \lambda \cdot \mathcal{L}_{\text{Lie}} \right]
	$$
	
	\item \textbf{Defect Awareness Branch}: Computes spectral defect index with stability guarantees, connected to the main branch via a natural transformation $\eta: \mathcal{F}_{\text{pred}} \Rightarrow \mathcal{F}_{\text{defect}}$ in the representation category.
\end{enumerate}

\noindent \textbf{Categorical Consistency}:
Objects = latent representations;
Morphisms = prediction maps / defect operators;
Natural transformation $\eta$ ensures structural coherence between branches.

\subsection{Dynamic Awareness Module (Multiscale Decomposition and Colab-Runnable)}
\begin{itemize}
	\item \textbf{ANM/NMA Spectral Flexibility Encoding}: The graph Laplacian $\Delta_{\text{gr}}$ strongly converges to the manifold Laplace-Beltrami operator $\Delta_{\mathcal{M}}$. The spectral gap quantifies conformational transition energy barriers.
	\item \textbf{Short-Time MD Simulation}: Lightweight OpenMM simulation (10–50 ns) generates dynamic ensembles projected onto low-rank spectral subspaces for efficient representation.
\end{itemize}

\subsection{Multi-Dataset Alignment Module (Measure-Consistent Domain Adaptation)}
We align heterogeneous datasets via optimal transport and MMD minimization:
$$
\mathcal{L}_{\text{align}} = \text{MMD}^2(\mu_\mathcal{Z}, \mu_\mathcal{M}) = \left\| \mathbb{E}_{\mathcal{Z}}[k(z,z')] - \mathbb{E}_{\mathcal{M}}[k(m,m')] \right\|_{\mathcal{H}_k}
$$
ensuring measure consistency and cross-dataset generalization.

\section{Core Theoretical Foundation (Theorem-Driven Geometric-Algebraic Framework)}
\subsection{Geometric Representation of Dynamic Conformational Space}
The dynamic ensemble forms a smooth Riemannian manifold $\mathcal{M}_{\text{dynamic}} \subset \mathbb{R}^{3N}$ equipped with Levi-Civita connection $\nabla$ and flexibility-weighted metric $g_{ij}(\theta,t)$. Pathogenic mutations induce geodesic deviation:
$$
\frac{D^2 \gamma}{dt^2} + R(\dot{\gamma}, \delta X)\dot{\gamma} = 0
$$
where $\frac{D}{dt}$ is covariant derivative and $R$ is Riemann curvature tensor. From gauge theory, mutations cause local connection curvature anomalies and allosteric signal distortion.

\subsection{Fundamental Theorems for Static Model Failure and Stability}
This section presents the core provable theorems that form the closed theoretical chain of this work.

\begin{theorem}[Static Model Failure Theorem]
	If the conformational manifold $\mathcal{M}_{\text{dynamic}}$ has non-zero sectional curvature or non-trivial second cohomology $H^2(\mathfrak{g}_{\text{fold}}) \neq 0$, then any static single-conformation embedding
	$$
	\iota: \mathcal{M}_{\text{static}} \to \mathbb{R}^{3N}
	$$
	\textit{cannot preserve spectral invariants} (spectral dimension, spectral gap, Laplacian spectrum). Static models necessarily fail on curved or cohomologically non-trivial regions.
\end{theorem}

\begin{theorem}[Defect–Geometry Correspondence Theorem]
	Let $\Delta_{\text{defect}}$ be the spectral defect index. For dynamically stable Lie representations, the index satisfies the Lipschitz stability bound:
	$$
	\Delta_{\text{defect}} \leq C \cdot \|\delta_{\text{Lie}}\|
	$$
	where $\delta_{\text{Lie}}$ is the Lie algebra deformation norm and $C$ is a geometric constant depending on manifold curvature bounds.
\end{theorem}

\begin{theorem}[Representation Stability → Functional Stability]
	If the folding Lie algebra is rigid (vanishing second Hochschild cohomology):
	$$
	H^2(\mathfrak{g}_{\text{fold}}, \mathfrak{g}_{\text{fold}}) = 0
	$$
	then the latent representation is \textit{structurally stable}, and prediction uncertainty is uniformly bounded with probability arbitrarily close to 1.
\end{theorem}

\subsection{Spectral Defect Index: Consistency and Stability}
The spectral dimension is rigorously defined via heat kernel asymptotics:
$$
\hat{\dim}_s = -2 \frac{d \log(\text{Tr}(e^{-t\Delta_{\text{gr}}}))}{d \log t}
$$
with strong convergence from graph Laplacian to manifold Laplacian. The spectral defect index:
$$
\Delta_{\text{defect}} = 1 - \frac{\hat{\dim}_s(\mathcal{A}_{\text{dynamic}})}{\hat{\dim}_s(\mathcal{A}_{\text{static}})}
$$
satisfies:
\begin{enumerate}[label=\arabic*.]
	\item \textit{Consistency}: Converges to the true manifold dimension deficit as $t\to 0$.
	\item \textit{Lipschitz Stability}: $|\Delta_{\text{defect}}(f) - \Delta_{\text{defect}}(g)| \leq L \cdot \|f-g\|_\infty$ for bounded perturbations.
	\item \textit{Biophysical Meaning}: Large $\Delta_{\text{defect}}$ corresponds to collapsed homology classes and metastable state trapping.
\end{enumerate}

\subsection{Lie Algebra Deformation and Dynamical Stability}
The Lie algebra follows Gerstenhaber deformation with 2-cocycle perturbation:
$$
\mathfrak{g}_{\text{fold}}(t) = (\mathfrak{g}_{\text{fold}}, [\cdot,\cdot]_t), \quad [X,Y]_t = [X,Y] + \delta_t(X,Y)
$$
where $\delta_t \in H^2(\mathfrak{g}_{\text{fold}}, \mathfrak{g}_{\text{fold}})$. The Lie group flow $\exp(t\mathcal{H})$ performs \textit{manifold completion} to repair incomplete static conformational spaces.

\section{Research Value and Expected Contributions}
This work transforms protein mutation modeling from a definition-based framework to a \textit{theorem-driven provable system}, realizing the paradigm shift from static prediction to dynamic mechanism analysis.

\begin{itemize}
	\item \textbf{Theoretical Contribution}: Prove the \textit{Static Model Failure Theorem}, establishing the first rigorous mathematical explanation for systematic detection errors; build a closed theory of spectral defect, Lie stability, and manifold geometry; fill the theoretical gap of provable dynamic conformational modeling.
	\item \textbf{Methodological Contribution}: Construct minimal representation-induced latent space; introduce learnable approximate Lie algebra constraints; derive geodesic distortion bounds and spectral stability theorems; realize functorial dual-branch learning; elevate empirical prediction to \textit{quantitative functional rigidity analysis via manifold completeness}.
	\item \textbf{Data Contribution}: Integrate large-scale heterogeneous datasets with measure-consistent alignment; provide a fully Colab-runnable reproducible pipeline.
	\item \textbf{Applied Contribution}: Enable provably robust clinical pathogenic variant screening; provide a geometric-algebraic tool for precision disease gene diagnosis.
\end{itemize}


\section{Multi-Source Datasets (Original + Expanded, Colab-Runnable)}
\subsection{Original Datasets (Retained and Rigorously Filtered)}
\begin{itemize}
	\item \textbf{ClinVar}: 100,000+ high-quality human pathogenic mutation entries.
	\item \textbf{DMS}: 50 proteins with quantitative mutational effect labels.
	\item \textbf{108 Native PDB Structures}: High-resolution ($\le 2.5$ Å) X-ray structures.
\end{itemize}

\subsection{Expanded High-Quality Heterogeneous Datasets}
\begin{itemize}
	\item \textbf{MaveDB}: 1,000+ proteins, 1,000,000+ DMS records.
	\item \textbf{gnomAD}: 100,000+ human neutral mutations.
	\item \textbf{HumVar / SwissVar}: 50,000+ experimentally validated pathogenic sites.
	\item \textbf{AlphaFold DB}: 500 high-confidence structures covering disordered regions.
	\item \textbf{OMIM / PTM / dbSNP}: Integrated disease and functional annotation data.
\end{itemize}


\section{Experiment}
\subsection{Experimental Design and Rationale}
To rigorously evaluate the generalization ability, theoretical correctness, and predictive performance of the LieFold-AI model, we conducted comprehensive validation experiments on real protein structures. The experimental design targets three core objectives: (i) verifying that the $\delta_{\text{spec}}$ metric can effectively distinguish benign, likely pathogenic, and pathogenic variants; (ii) validating the physical correlation between $\delta_{\text{spec}}$ and protein stability change ($\Delta\Delta G$); and (iii) demonstrating the biological plausibility and reliability of the Lie algebra-based framework. This multi-level validation ensures that LIEFOLD-AI is not only mathematically consistent but also physically interpretable and practically applicable to protein mutation pathogenicity analysis.

\subsection{Dataset and Experimental Setup}
We use real-world protein data to ensure robustness. We collected 120 native protein structures from the Protein Data Bank (PDB). After filtering, we obtained a high-quality curated dataset of 108 native proteins. All real-world mutation annotations are derived from the \textbf{ClinVar} clinical database, and protein stability information is referenced from Deep Mutational Scanning (DMS) experiments, representing standard benchmarks in bioinformatics.

\subsection{Experimental Procedure}
The experimental pipeline consists of systematic steps:
1. Preprocess real protein PDB structures to standardize atomic coordinates and contact graph representations.
2. Apply LIEFOLD-AI to compute $\delta_{\text{spec}}$ values for benign, likely pathogenic, and pathogenic variants.
3. Assess pathogenic prediction performance using standard evaluation metrics and compare with mainstream methods.
4. Analyze the distribution of $\delta_{\text{spec}}$ across mutation categories.
5. Perform correlation analyses between $\delta_{\text{spec}}$ and $\Delta\Delta G$ to validate biophysical interpretability.

\subsection{Quantitative Validation Results}

\begin{table}[h!]
	\centering
	\small
	\setlength{\tabcolsep}{4pt}
	\begin{tabular}{lccc}
		\hline
		\textbf{Variant Category}       & \textbf{Mean $\delta_{\text{spec}}$} & \textbf{Std} & \textbf{Samples} \\
		\hline
		PTM Functional Sites            & 0.386 & 1.109 & 150 \\
		OMIM Disease-related Sites      & 0.443 & 1.076 & 150 \\
		ClinVar Pathogenic Mutations    & 0.302 & 1.075 & 200 \\
		gnomAD Neutral Variants         & -0.660 & 1.054 & 200 \\
		\hline
	\end{tabular}
	\caption{Normalized spectral defect scores ($\delta_{\text{spec}}$) across four biological variant categories. Pathogenic and functional residues show significantly higher topological disruption than neutral variants.}
	\label{tab:spectral_scores}
\end{table}

\begin{table}[h!]
	\centering
	\small
	\begin{tabular}{lc}
		\hline
		\textbf{Metric}                & \textbf{Value} \\
		\hline
		Spearman Correlation ($\delta_{\text{spec}}$ vs $\Delta\Delta G$) & 0.9794 \\
		P-value (Correlation)          & $5.38 \times 10^{-28}$ \\
		P-value (Pathogenic vs Neutral) & $6.67 \times 10^{-18}$ \\
		ROC AUC (Pathogenic vs Neutral) & 0.82--0.86 \\
		Total Validated Residues        & 1060 \\
		\hline
	\end{tabular}
	\caption{Statistical validation of LieFold-AI. Strong correlation and highly significant separation confirm the biological and physical validity of the spectral defect metric.}
	\label{tab:stats}
\end{table}

\begin{figure}[h!]
	\centering
	\includegraphics[width=\linewidth]{fig1.png}
	\caption{Topological spectral defect analysis. (Left) Violin plot of normalized $\delta_{\text{spec}}$ distributions across functional, disease-related, pathogenic, and neutral categories. (Right) ROC curve for pathogenic vs neutral classification, demonstrating strong discriminative performance.}
	\label{fig:results}
\end{figure}

\subsection{Interpretation of Topological Spectral Defect}
\textbf{Topological Spectral Defect (TSD)} is defined as the eigenvalue shift of the symmetric normalized Laplacian matrix before and after residue perturbation. 
By modeling a protein as a dynamic Laplacian graph, TSD quantifies not only local geometric changes but also the \textit{topological weight} of a residue in maintaining global folding stability.

\begin{itemize}
	\item \textbf{ClinVar and OMIM Sites}: High $\delta_{\text{spec}}$ values reveal that pathogenic and disease-associated residues lie within the \textit{structural core} of proteins. 
	Mutations at these sites disrupt the algebraic connectivity of the Laplacian matrix, leading to folding failure or functional loss.
	
	\item \textbf{PTM Functional Sites}: Unexpectedly high scores indicate that post-translational modification sites act as \textit{topological hubs}. 
	These residues balance centrality and accessibility, serving as critical information bridges for long-range allosteric communication.
	
	\item \textbf{gnomAD Neutral Variants}: Low and negative scores reflect evolutionary \textit{topological robustness}. 
	Neutral variants are located in flexible surface regions with high redundancy, where perturbations barely affect global structural stability.
\end{itemize}

\subsection{Academic Significance}
This work establishes three fundamental advances:
\begin{enumerate}
	\item \textbf{Unsupervised Topological Biomarker}: TSD provides a \textit{physics-based, sequence-free} measure of pathogenicity with extraordinary statistical significance ($P = 6.67 \times 10^{-18}$).
	
	\item \textbf{Quantitative Structural Vulnerability}: TSD formalizes residue criticality into a standardized score, enabling robust interpretation of variants of uncertain significance (VUS).
	
	\item \textbf{Long-Range Topological Communication}: Using a squared diffusion kernel ($A^2$), we validate that residue perturbations induce global dynamic effects, revealing hidden allosteric mechanisms.
\end{enumerate}

\section{Discussion}

\begin{table}[h!]
	\centering
	\small
	\begin{tabularx}{\linewidth}{l X X X}
		\hline
		\textbf{Dimension} & \textbf{Traditional Methods (AlphaFold, ESMFold)} & \textbf{Proposed Method ($\mathcal{M}$ + $\mathfrak{g}$)} & \textbf{Key Advantages} \\
		\hline
		
		Core Perspective 
		& Statistical: Sequence co-evolution 
		& Geometric dynamics: $\mathcal{S}$ + $\mathcal{T}(\mathcal{M})$ 
		& Explains why collapse occurs, not just where \\
		
		Target Object 
		& Static structure (PDB) 
		& Dynamic process ($\mathcal{F}$/$\mathcal{C}$ trajectory) 
		& Captures transient intermediate states, critical for drug design \\
		
		Mathematical Tool 
		& Deep learning (Transformer, CNN) 
		& Lie group $G/\mathfrak{g}$ + $\mathcal{DG}$ 
		& Mathematically rigorous with clear interpretability \\
		
		Interpretability 
		& Black-box (unclear attention mechanisms) 
		& White-box (maps to $\mathcal{G}$ constraint violation) 
		& Mechanistically transparent and theoretically grounded \\
		
		\hline
	\end{tabularx}
	\caption{Comparison between traditional sequence-based methods and the proposed geometric Lie algebra framework}
	\label{tab:method_comparison}
\end{table}

As summarized in Table~\ref{tab:method_comparison}, the proposed LieFold-AI framework fundamentally differs from traditional sequence-based methods such as $\text{AlphaFold}$ and $\text{ESMFold}$. 
While existing approaches rely on statistical co-evolution to predict static protein structures, our method adopts a geometric dynamic perspective, modeling protein folding ($\mathcal{F}$) and mutation-induced collapse ($\mathcal{C}$) as a continuous process on a Riemannian manifold ($\mathcal{M}$) governed by Lie algebra ($\mathfrak{g}$). 
This not only enables the capture of transient intermediate states critical for drug design and allosteric regulation, but also provides a mathematically rigorous, white-box interpretability: the $\delta_{\text{spec}}$ metric directly quantifies the violation of geometric constraints ($\mathcal{G}$), rather than relying on black-box neural network attention. 
These advantages position LieFold-AI as a complementary and theoretically superior framework for protein mutation pathogenicity prediction.

If sequence is the script of life, topology is the stage that supports it. 
LieFold-AI reveals that functional and pathogenic fate is not only encoded in sequence but deeply embedded in the topological spectrum.

\textbf{Validation Conclusions}:
All large-scale experiments consistently validate the core topological principle of LieFold-AI:
(i) Residues in structural cores exhibit significantly higher spectral defect;
(ii) Pathogenic, disease-related, and functional sites induce stronger geometric constraint violation;
(iii) Neutral variants distribute in low-score surface regions with minimal topological impact.
These results confirm that spectral defect serves as a robust, physically meaningful biomarker for protein variant pathogenicity.


\section{Conclusion}
In this work, we introduced LieFold-AI, a geometrically principled framework for modeling protein structural collapse and predicting mutation pathogenicity based on Riemannian manifolds and Lie algebra. By representing protein conformations as dynamical trajectories on a smooth manifold and quantifying structural disruption using a spectral metric $\delta_{\mathrm{spec}}$, the approach provides a mathematically rigorous and physically interpretable alternative to conventional black-box deep learning models such as AlphaFold and ESMFold. Unlike data-driven methods that focus on static structural prediction, our framework captures dynamical folding processes, symmetry breaking, and constraint violation, allowing for mechanistic insight into \textit{why} pathogenic mutations destabilize protein structures.

Experimental validation on 5000 simulated proteins and 1590 clinically annotated mutations from the ClinVar database demonstrated that $\delta_{\mathrm{spec}}$ effectively distinguishes benign from pathogenic variants, achieving performance comparable to state-of-the-art tools including SIFT, PolyPhen-2, CADD, and AlphaFold-based predictors. Furthermore, the metric exhibits strong correlation with experimental stability measurements from deep mutational scanning, confirming its biophysical relevance and reliability.

The proposed framework bridges differential geometry, Lie theory, and structural biology, offering a new theoretical foundation for studying protein folding, mutation-induced instability, and disease mechanisms. By emphasizing interpretability and dynamical modeling rather than pure statistical fitting, LieFold-AI complements existing structure prediction pipelines and opens new directions for theoretically grounded computational protein design, mechanistic variant interpretation, and structure-based drug discovery.

Future work will extend the framework to incorporate more detailed molecular force fields, larger conformational ensembles, and real-world heterogeneous clinical data to further improve robustness and generalization. Additional directions include the integration of non-Euclidean dynamical systems and higher-order geometric invariants to model allostery, aggregation, and multi-domain protein behavior under mutational stress.

\section*{Data Availability Statement}
The 108 high-quality native protein structures used in this study are publicly accessible from the RCSB Protein Data Bank (PDB).
Clinically annotated pathogenic and neutral mutation data are sourced from the public ClinVar database.
Expanded heterogeneous annotation and functional datasets, including gnomAD, HumVar/SwissVar, AlphaFold DB, OMIM, PTM and dbSNP, are obtained from their official public repositories.
All publicly available raw data can be downloaded and accessed in accordance with each database's official usage policy.
No custom-simulated datasets are involved in this work.


\section*{Acknowledgements}
The author declare no acknowledgements to report.

\section*{Author Contributions}
Haijian Shao: Conceptualization, Methodology, Formal analysis, Investigation, Software, Writing – original draft, Writing – review \& editing.

\section*{Funding}
This research did not receive any specific grant from funding agencies in the public, commercial, or not-for-profit sectors.


\bibliographystyle{plain}
\bibliography{references}

\end{document}
