## Supplementary material for "Geometric Theoretical Framework for Dynamic Protein Mutation Detection Models: Defect Awareness and Pathogenicity Prediction": main latex: references.bib

@article{begun2025local,
  title={Local topology and perestroikas in protein structure and folding dynamics},
  author={Begun, Alexander and Chernodub, Maxim N and Molochkov, Alexander and Niemi, Antti J},
  journal={Physical Review E},
  volume={111},
  number={2},
  pages={024406},
  year={2025},
  publisher={APS}
}

@article{wei2020protein,
  title={Protein structure determination using a Riemannian approach},
  author={Wei, Xian and Li, Zhi-Cheng and Li, Shi-Jian and Peng, Xu-Biao and Zhao, Qing},
  journal={FEBS letters},
  volume={594},
  number={6},
  pages={1036--1051},
  year={2020},
  publisher={Wiley Online Library}
}

@article{diepeveen2024riemannian,
  title={Riemannian geometry for efficient analysis of protein dynamics data},
  author={Diepeveen, Willem and Esteve-Yagüe, Carlos and Lellmann, Jan and Öktem, Ozan and Schönlieb, Carola-Bibiane},
  journal={Proceedings of the National Academy of Sciences},
  volume={121},
  number={33},
  pages={e2318951121},
  year={2024},
  publisher={National Academy of Sciences}
}

@article{jumper2021highly,
  title={Highly accurate protein structure prediction with AlphaFold},
  author={Jumper, John and Evans, Richard and Pritzel, Alexander and Green, Tim and Figurnov, Michael and Ronneberger, Olaf and Tunyasuvunakool, Kathryn and Bates, Russ and Žídek, Augustin and Potapenko, Anna and others},
  journal={nature},
  volume={596},
  number={7873},
  pages={583--589},
  year={2021},
  publisher={Nature Publishing Group UK London}
}

@article{isert2023structure,
  title={Structure-based drug design with geometric deep learning},
  author={Isert, Clemens and Atz, Kenneth and Schneider, Gisbert},
  journal={Current Opinion in Structural Biology},
  volume={79},
  pages={102548},
  year={2023},
  publisher={Elsevier}
}

@article{banerjee2025accurate,
  title={Accurate identification and mechanistic evaluation of pathogenic missense variants with Rhapsody-2},
  author={Banerjee, Anupam and Bogetti, Anthony T and Bahar, Ivet},
  journal={Proceedings of the National Academy of Sciences},
  volume={122},
  number={18},
  pages={e2418100122},
  year={2025},
  publisher={National Academy of Sciences}
}

@article{li2024prediction,
  title={Prediction of mutation-induced protein stability changes based on the geometric representations learned by a self-supervised method},
  author={Li, Shan Shan and Liu, Zhao Ming and Li, Jiao and Ma, Yi Bo and Dong, Ze Yuan and Hou, Jun Wei and Shen, Fu Jie and Wang, Wei Bu and Li, Qi Ming and Su, Ji Guo},
  journal={BMC bioinformatics},
  volume={25},
  number={1},
  pages={282},
  year={2024},
  publisher={Springer}
}

@inproceedings{atiyah1975spectral,
  title={Spectral asymmetry and Riemannian geometry. I},
  author={Atiyah, Michael F and Patodi, Vijay K and Singer, Isadore M},
  booktitle={Mathematical Proceedings of the Cambridge Philosophical Society},
  volume={77},
  number={1},
  pages={43--69},
  year={1975},
  organization={Cambridge University Press}
}

@article{walther2007signatures,
  title={Signatures of hydrophobic collapse in extended proteins captured with force spectroscopy},
  author={Walther, Kirstin A and Gräter, Frauke and Dougan, Lorna and Badilla, Carmen L and Berne, Bruce J and Fernandez, Julio M},
  journal={Proceedings of the National Academy of Sciences},
  volume={104},
  number={19},
  pages={7916--7921},
  year={2007},
  publisher={National Academy of Sciences}
}
