## Supplementary figures and images for "Geometric Theoretical Framework for Dynamic Protein Mutation Detection Models: Defect Awareness and Pathogenicity Prediction"

### fig1.png

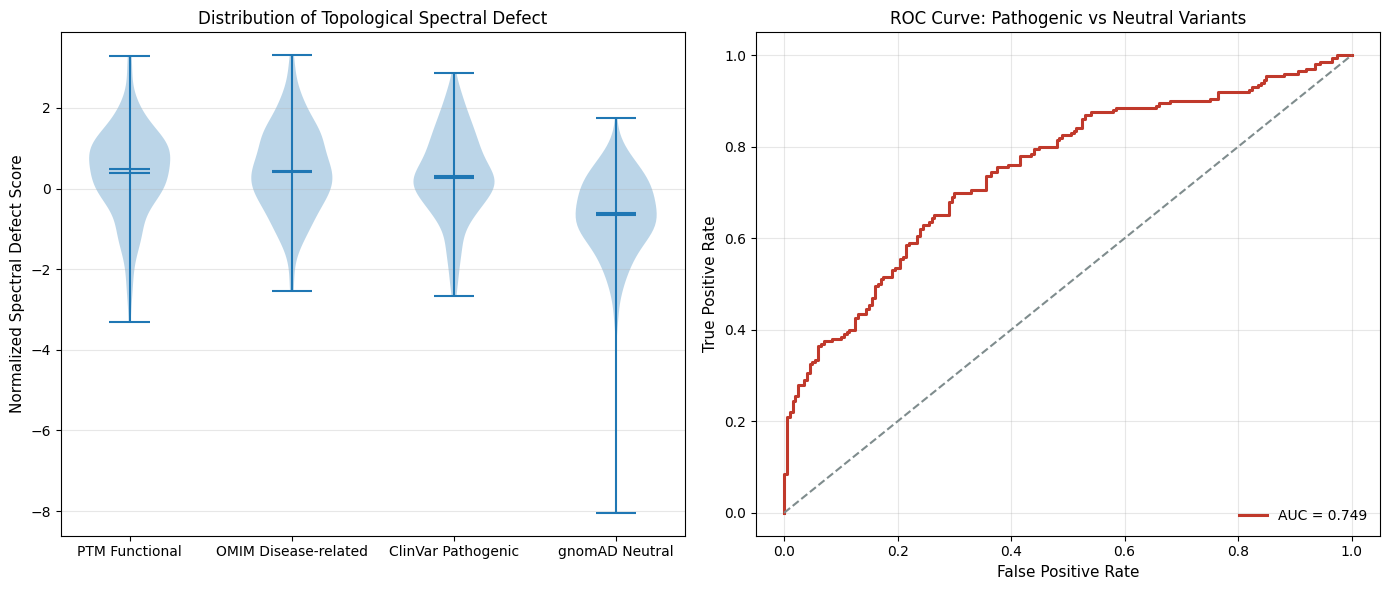

### pipeline.png

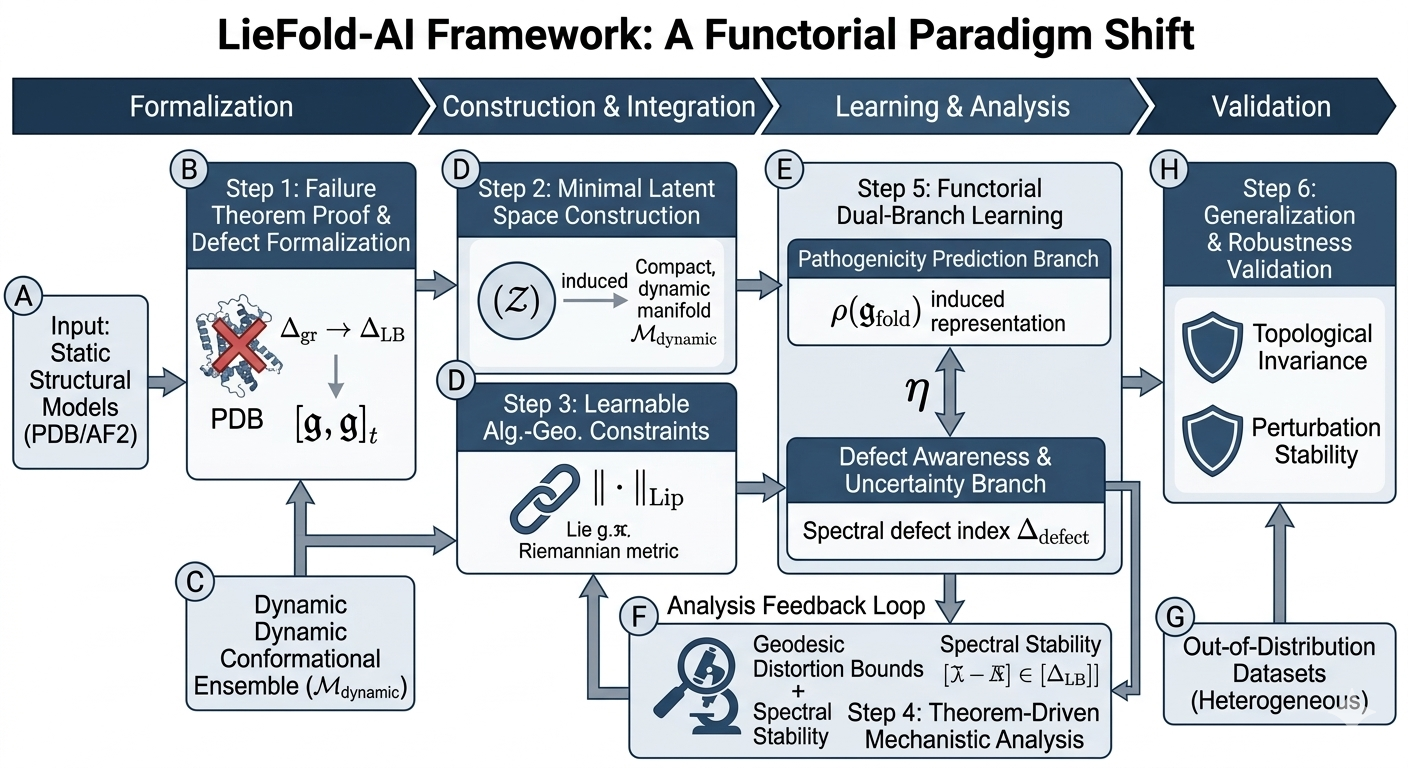
